# A high-throughput approach reveals distinct peptide charging behaviors in electrospray ionization mass spectrometry

**DOI:** 10.1101/2023.03.31.535171

**Authors:** Allyn M. Xu, Lauren C. Tang, Marko Jovanovic, Oded Regev

## Abstract

Electrospray ionization is a powerful and prevalent technique used to ionize analytes in mass spectrometry. The distribution of charges that an analyte receives (charge state distribution, CSD) is an important consideration for interpreting mass spectra. However, due to an incomplete understanding of the ionization mechanism, the analyte properties that influence CSDs are not fully understood. Here, we employ a machine learning-based high-throughput approach and analyze CSDs of hundreds of thousands of peptides. Interestingly, half of the peptides exhibit charges that differ from what one would naively expect (number of basic sites). We find that these peptides can be classified into two regimes—undercharging and overcharging—and that these two regimes display markedly different charging characteristics. Strikingly, peptides in the overcharging regime show minimal dependence on basic site count, and more generally, the two regimes exhibit distinct sequence determinants. These findings highlight the rich ionization behavior of peptides and the potential of CSDs for enhancing peptide identification.

## Introduction

Over the years, electrospray ionization^1^ (ESI) has become a leading ionization technique for pairing with liquid chromatography tandem mass spectrometry (LC-MS/MS)^2, 3^. ESI’s ability to ionize a wide range of biomolecules, and to process samples in a high-throughput manner^4–8^ has greatly broadened the scope of mass spectrometry^9, 10^, enabling applications to proteomics^11–14^, clinical biology^15, 16^, drug discovery^17^, and more^18^.

ESI ionizes aqueous solutions through maintaining a high voltage potential across a capillary, which vaporizes the solution into a mist of highly charged droplets^19^. As the solvent continues to evaporate, the droplets experience an increased charge density, and fissure into smaller droplets upon reaching their Rayleigh limit^20^, the point at which the Coulombic forces overcome the surface tension. The charges are ultimately deposited onto the analytes, through mechanisms that are still not fully understood^3, 21^, producing gaseous ionic molecules that arrive at the mass analyzer. Several theories have been proposed to explain the ionization mechanism, such as the ion evaporation model^22, 23^, the charge residue model^24, 25^, and the chain-ejection model^26, 27^, which differ in their assumptions of how the analyte interacts with the droplet. It is believed that these evaporation models are appropriate for different types of analytes, depending on their size and structure^27^. However, the dynamic nature of ESI and the inability to directly observe ESI at the molecular scale have made it challenging to fully characterize the determinants of analyte ionization.

Here we utilize a high-throughput approach to investigate the ESI ionization of peptides. We reasoned that a systematic analysis of a large-scale dataset would not only complement existing studies on select analytes, but also provide more insights than previous “black box”^28^ deep learning approaches^29^. We therefore generated a dataset containing charging information on hundreds of thousands of peptides, using both new and published LC-MS/MS runs. For each peptide, the resulting dataset includes the measured charge state distribution (CSD), defined as the relative intensities of the ions produced by that peptide.

We next employed machine learning on this dataset to gain insights into the relationship between peptide sequence and CSD. Our analysis revealed that half of the peptides exhibited charges that did not correspond to their number of basic sites. Classifying these peptides into two regimes, namely undercharging and overcharging, we identified striking differences in their ionization characteristics. Specifically, we discovered that for overcharged peptides, mass takes precedence over basic site count, and that charging in the two regimes is affected by distinct amino acid features.

Overall, our findings offer new insights into the complex dynamics of peptide ionization, highlight that CSDs contain rich information about peptide sequences, and may open opportunities for applications to identification pipelines in proteomics.

## Results

### Overview of peptide CSD dataset

To facilitate a machine learning approach, we developed an extraction scheme to extract CSD readings from MS1 scans (see methods). In each LC-MS/MS run, a single CSD reading was assigned to each MS2-identified peptide by averaging CSD readings across the peptide’s elution.

To cover a wide range of experimental settings, we applied our extraction scheme to 326 positive-ion mode LC-MS/MS runs acquired from three sources: our own, Confetti^30^, and Meier et al.^31^ These data sources differ in their choice of experimental parameters, protease, organism, and type of mass spectrometry instrument (Orbitrap^32^ and timsTOF^33, 34^). The resulting dataset contained CSD readings of 261,667 unique peptides (Table 1).

**Table 1.** Overview of peptide CSD dataset. Breakdown of extracted CSDs from each of the three data sources (ours, Confetti, Meier et al.). Asterisk (*) indicates the number of LC-MS/MS runs after aggregating over fractionations.

|  | Data source |  |  | Total |
| --- | --- | --- | --- | --- |
|  | Ours | Confetti | Meier et al. |  |
| LC-MS/MS runs | 20 | 18 | 288 (39*) | 326 (77*) |
| Total peptides | 264,259 | 166,543 | 416,306 | 847,108 |
| Unique peptides | 41,594 | 80,402 | 183,340 | 261,667 |
| Varied parameters | gradient length,<br>voltage, flow rate | protease | organism,<br>protease |  |
| MS instrument | Orbitrap | Orbitrap | timsTOF |  |

In order to confirm the reproducibility of the extracted CSD readings, we compared CSDs obtained from various LC-MS/MS runs. We found that CSD readings of the same peptide were generally consistent across different runs (Supplementary Fig. 1), and especially consistent among experimental replicates (*∼*3.7% error Orbitrap, *∼*5.2% error timsTOF). After applying a one-parameter batch correction (see methods), errors across replicates dropped slightly (*∼*3.4% error Orbitrap, *∼*4.7% timsTOF). Batch correction was not used in downstream analysis as LC-MS/MS runs were analyzed separately. Furthermore, the datasets from timsTOF instruments had slightly higher errors (Supplementary Fig. 1), which may be due to the extraction scheme being insufficiently optimized to that technology (as it does not employ ion mobility information). Together, these results demonstrate the robustness of the extraction scheme, and that peptide CSDs are highly consistent across replicates.

Peptides in our dataset exhibited a variety of CSDs (select peptides shown in Fig. 1). It is known that the basic site count serves as a rough estimate for the charge a peptide receives in positive-ion mode ESI^35^. Indeed, 51% of the peptides in the dataset have a CSD concentrated solely on the charge state equal to the basic site count. On the other hand, 40% of peptides exhibit undercharging (mean charge less than basic site count), and 9% exhibit overcharging (mean charge greater than basic site count). In downstream analysis, we explore factors that explain why some peptides receive fewer charges or more charges than their basic site count.

**Figure 1.**
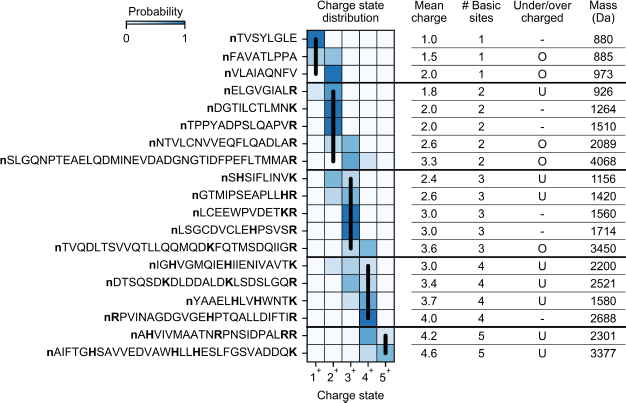
CSDs of select peptides. Extracted CSDs and properties of select peptides from one of our HeLa trypsin runs (2.5 kV ESI voltage, 160 min gradient length, 400 nL/min flow rate). Basic sites are bolded. Vertical black lines denote the charge state equal to basic site count. U = undercharged (mean charge *<* basic site count). O = overcharged (mean charge *>* basic site count).

### Under- and overcharged peptides exhibit different dependence on mass and number of basic sites

Grouping peptides by their basic site count, we observed that mean charge increases with mass, and exhibits transitions from under- to overcharging in all LC-MS/MS runs (representative runs shown in Fig. 2, Supplementary Figs. 3, 4). For instance, in one of our HeLa runs, among peptides with three basic sites and mass less than 2600 Da, 98% exhibited charges of 3^+^ or lower (undercharging, Fig. 2b). In contrast, among peptides with three basic sites and mass greater than 2600 Da, 94% exhibited charges 3^+^ or higher (overcharging, Fig. 2b). We confirmed that this transition phenomenon cannot be explained by the mass spectrometer’s *m*/*z* cutoff, since the presence of a charge state recedes well before reaching the *m*/*z* cutoff. Moreover, the phenomenon is present in runs performed with other proteases (Supplementary Fig. 3) and with the timsTOF instrument (Supplementary Fig. 4), suggesting it is independent of the peptide distribution or the choice of mass spectrometer instrument.

**Figure 2.**
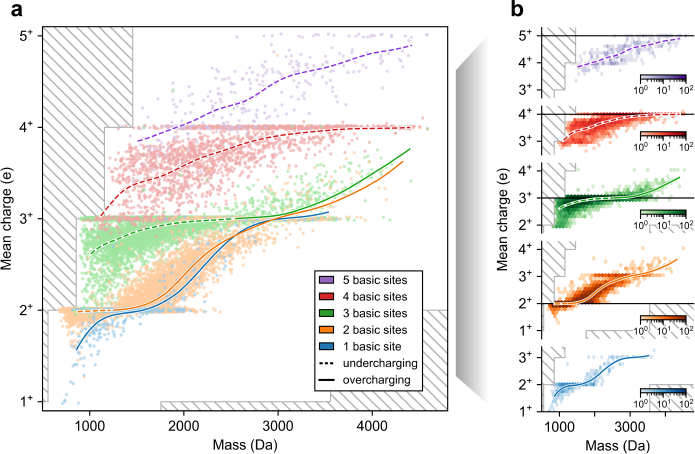
Visualizing CSD dataset versus mass. Plots of mean charge versus mass for peptides from a representative run (our HeLa trypsin run; 2.5 kV ESI voltage, 160 min gradient length, 400 nL/min flow rate), colored by number of basic sites. Data points are shown as (a) a scatter plot with jittering in the y-axis, uniformly chosen from −0.02 to 0.02, and (b) 2D hexagonal binning plots separated by basic site count. Colored curves show spline-interpolation of mean charge versus mass, performed with 20 cubic-splines, a smoothing parameter of 5, and a monotonicity constraint. Curves are dashed when the interpolated mean charge is less than the basic site count (undercharging) and solid otherwise (overcharging). Hatched regions correspond to unobservable mean charge due to *m*/*z* cutoff.

**Figure 3.**
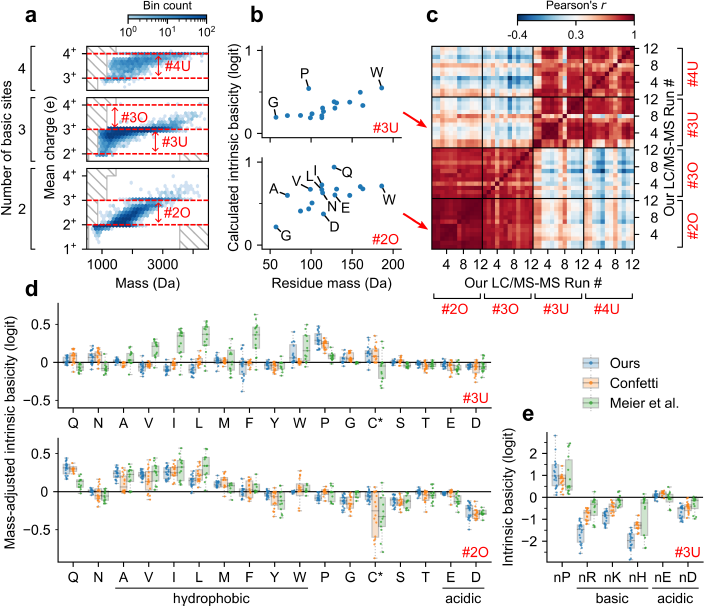
CSD sequence determinants for under- and overcharged peptides. (a) Schematic representation of the four peptide charging regions of interest, determined by number of basic sites and pair of consecutive charge states (see text). (b) Scatter plots of calculated amino acid intrinsic basicities (see text) versus mass for two representative charging regions (top: region #3U; bottom: region #2O). Panels (a–b) use our HeLa trypsin run with 2.5 kV ESI voltage, 160 min gradient length, 400 nL/min flow rate. (c) Heatmap showing correlations between calculated amino acid intrinsic basicities among all pairs of region-run combinations for our runs with sufficiently many data points (see methods). Along both axes, all 12 runs are aligned four times, once for each of the four regions. (d) Box plot of estimated mass-adjusted intrinsic basicities (given by residual intrinsic basicities after subtracting trend in mass, see methods) across runs with sufficiently many data points, separated by data source and charging region. Runs from Meier et al. were aggregated across fractionations (average taken for duplicate readings). C* denotes carbamidomethyl-cysteine in our and Meier et al.’s runs, and N-ethylmaleimide modified cysteine in Confetti’s runs. (e) Box plot of intrinsic basicities for the N-terminal identity of select amino acids (region #3U). Box plot elements: centerline, median; boxplot limits, upper and lower quartiles; whiskers, 1.5x interquartile range; points, in whisker range with jitters (outliers not shown).

Prior studies on select protein complexes^36–38^ and synthetic peptides^39^ have also observed correlations between size and charging. Moreover, the relationship is consistent with proposed ionization theories^27^ as larger analytes would be enclosed in larger, more highly charged droplets^20^, may have higher solvent-exposed surface area^37, 38^, and may have more functional groups that can stabilize charge through intramolecular solvation^40–42^. Our findings suggest that this correlation holds universally for peptides, and provide quantitative dependencies between the number of basic sites, mass, and mean charge.

One striking observation is that in the overcharging regime, the number of basic sites has little effect on the mean charge trend as a function of mass (Fig. 2a, solid lines). In contrast, in the undercharging regime, the number of basic sites has a significant effect on the mean charge trend (Fig. 2a, dashed lines). Thus, in the overcharging (but not in the undercharging) regime, mass takes precedence over the number of basic sites. For instance, among peptides of mass 1800 Da, those with 1 or 2 basic sites exhibit, on average, similar mean charges of 2.1 and 2.2, respectively (overcharging), whereas those with 3, 4, or 5 basic sites exhibit, on average, distinct mean charges of 2.9, 3.4, and 4.0, respectively (undercharging). These observations suggest differences in ionization factors at play for the over- and undercharging regimes.

These findings also demonstrate the potential benefits that CSDs can offer to peptide identification. Aside from peptides that exhibit overcharging, peptides sharing the same mass but having a different number of basic sites generally exhibit different CSDs (Supplementary Fig. 5), allowing them to be distinguished based on the MS1 scan. For example, among peptides with mass 1600 *±* 25 Da, 99% of those with two basic sites have mean charge < 2.5, while 98% of those with three basic sites have mean charge > 2.5. As such, when one is searching for potential peptide candidates for a given collection of ion peaks, the observed mass and CSD can be used to infer the peptide’s basic site count. This example showcases a preliminary use-case for peptide CSDs, and suggests that incorporating more sequence-dependent insights may provide further identification opportunities.

### Distinct sequence determinants underlie peptide ionization in the under- and overcharg-ing regimes

A peptide’s ionization depends on many factors, including Coulombic forces^43, 44^, its gas-phase conformation^41, 45^, its protonatable locations^46, 47^, and intramolecular forces^40–42^. To provide further insights into the sequence factors that influence peptide ionization, we analyzed the sequence determinants of CSDs within the under- and overcharging regimes.

For the analysis, we considered the 20 amino acid counts and the identity of the N-terminal amino acid as the sequence features of interest. To account for possible dependence on the number of basic sites or charge states, we performed a separate analysis for each of four regions, labelled #2O, #3O, #3U, and #4U (O : overcharged, U : undercharged, Fig. 3a). Region #2O consists of peptides with two basic sites and considers charging across 2^+^ and 3^+^. We similarly define region #3O (three basic sites, charging across 3^+^ and 4^+^), region #3U (three basic sites, charging across 2^+^ to 3^+^), and region #4U (four basic sites, charging across 3^+^ and 4^+^). For each of our 12 LC-MS/MS runs (with sufficiently many data points, see methods) and each of the four regions, we calculated a feature’s contribution using an “intrinsic basicity” score (Fig. 3b), derived from coefficients of a logistic regression (see methods). Interestingly, clustering the region-run pairs by their calculated intrinsic basicities, we identified two distinct clusters (Fig. 3c): there was strong agreement only between regions #2O and #3O, and between regions #3U and #4U. This result suggests that under- and overcharging are influenced by different sequence features, and that these sequence features do not depend strongly on the number of basic sites.

To understand the sequence determinants of overcharging, we examined some of the more prominent calculated intrinsic basicities for region #2O across all runs. We observed that glutamine (Q) and aspartic acid (D) have the highest and lowest mass-adjusted intrinsic basicities, respectively (Fig. 3d, bottom; Supplementary Fig. 6a, bottom). These observations are consistent with previous work: glutamine (Q) has been reported to have high gas-phase basicity due to its amide group^46, 48^ and can be a potential charge carrier during ESI^46^; aspartic acid (D) has been observed to form salt bridges with basic sites in molecular dynamics simulations^49^. Interestingly, despite having the same functional group, glutamine (Q) displays notably higher calculated intrinsic basicity than asparagine (N); similarly, glutamic acid (E) displays notably higher calculated intrinsic basicity than aspartic acid (D) (Fig. 3b,d). Lastly, we observed that amino acids with nonaromatic hydrocarbon side chains (alanine (A), valine (V), isoleucine (I), and leucine (L)) have high mass-adjusted intrinsic basicities.

Next, to understand the impact of sequence features on undercharging, we examined some of the more prominent calculated intrinsic basicities for region #3U across all runs. We observed that the presence of an arginine (R), lysine (K), or histidine (H) at the N-terminal greatly reduces charging (Fig. 3e; Supplementary Fig. 6b, top). This suggests that undercharging occurs due to Coulombic repulsion between the N-terminus and an N-terminal basic side chain. In particular, this effect can be observed in sequential isomers TNSTFNQVVLKR and RTNSTFNQVVLK, which have identical sequences apart from an arginine at the C- or N-terminal. In the four runs that contain CSD readings for both peptides, the former peptide has a CSD of (*p*_2+_ , *p*_3+_ ) = (4%, 96%) *±* 7% (mean *±* s.d.; not batch corrected), while the latter, with an N-terminal arginine, has a CSD of (*p*_2+_ , *p*_3+_ ) = (44%, 56%) *±* 6%. In addition to these Coulombic effects, we observed that proline (P) has a notably high intrinsic basicity (Fig. 3d, top; Supplementary Fig. 6a, top), especially if located at the N-terminal. Moreover, the runs from Meier et al. demonstrated higher intrinsic basicity for strongly hydrophobic amino acids (isoleucine (I), leucine (L), phenylalanine (F)) than did the Orbitrap runs (Fig. 3d, top).

In conclusion, our analysis demonstrates that the under- and overcharging regimes are influenced by distinct sequence determinants. These sequence determinants are consistent across different LC-MS/MS runs, and align with previously documented effects. Moreover, our findings indicate that peptide ionization depends on many factors beyond mass and basic site count, including both composition and position of the constituent amino acids. Collectively, these results underscore the value of CSDs in providing information about peptide sequences, and suggest that CSDs may complement other measured analyte properties for proteomics identification.

### CSD variations offer opportunities for identification

It is well known that certain LC-MS/MS experimental parameters change the overall degree of charging experienced by analytes^3, 35, 50, 51^. Indeed, in our own experiments, we observed that increasing flow rate and gradient length generally resulted in higher overall charging (Supplementary Fig. 7). While such experimental variations may be perceived as undesirable, they can be potentially harnessed to improve peptide identification. Specifically, peptides with similar CSDs may exhibit more distinct CSDs under different charging conditions, allowing for easier separation.

To illustrate this concept, consider the two peptides ASGQAFELILSPR and ACANPAAGSVILLENLR. In the low flow rate runs (200 nL/min), these peptides exhibited CSDs of (*p*_2+_ , *p*_3+_ ) = (96%, 4%) and (88%, 12%), respectively, a difference of 8%. In the high flow rate runs (800 nL/min), these peptides exhibited CSDs of (*p*_2+_ , *p*_3+_ ) = (90%, 10%) and (74%, 26%), respectively, a difference of 16%. In other words, increasing the flow rate resulted in a twofold increase in the difference between these two CSDs. In fact, this example occurs throughout our dataset (Supplementary Fig. 2a,b). For instance, among peptides with mean charge between 2 and 2.15 in the low flow rate run, more than half of the pairwise differences (59%) increased by twofold or more when flow rate increased (Supplementary Fig. 2a). Similarly, among peptides with mean charge between 2.85 and 3 in the high flow rate run, more than half of of the pairwise differences (52%) increased by two fold or more when flow rate decreased. Lastly, CSD variations across other pairs of runs exhibited similar trends (Supplementary Fig. 2b). Together, these findings highlight how varying experimental parameters can magnify differences in CSDs.

Instead of varying experimental parameters across runs, one can also imagine inducing variations in charging within a single run, possibly on a scan-to-scan basis. In fact, we observed that in many runs, mild scan-to-scan charging fluctuations already exist. While the reasons for these fluctuation are not clear, our findings indicate that they are not due to noise and have potential for improving identification (Supplementary Text 1).

In summary, these findings suggest that varying charging conditions across runs and within a single run may enhance the utility of CSDs for peptide identification.

## Discussion

In this work, we demonstrate that utilizing a high-throughput approach can reveal novel, general patterns underlying peptide ionization. Previous work has demonstrated that the ionization of native proteins, due to their tightly compacted structure in their folded state, is well-predicted by their surface-exposed area^37, 38^. In contrast, our study indicates that denatured peptides, typically encountered in shotgun proteomics^52^, exhibit complex ionization behavior: the interplay of mass and basic site count determines under- and overcharging, with further fine-tuning based on additional sequence-dependent factors.

Notably, our findings demonstrate that the under- and overcharging regimes display different dependencies on mass and basic site count. We found that basic site count has little effect on the typical mean charge exhibited by overcharged (but not undercharged) peptides. One potential explanation for this phenomenon is that overcharged peptides, having high mass relative to their basic site count, contain enough backbone carbonyls for solvating excess protons^42^. As such, additional basic sites may not have a strong effect on overcharging. There may be other potential explanations for this phenomenon, even causes arising outside of ESI such as the efficiency of transporting ions into the mass analyzer. Regardless, the striking contrast in basic site count dependence suggests there are differences in the underlying ionization processes that govern under- and overcharging of peptides.

Additionally, the distinct sequence determinants that we derived for the two regimes align with factors related to gas-phase interactions and dynamics, suggesting possible mechanistic connections. Specifically, for overcharged peptides, our findings align with previously documented effects of glutamine (Q) having high gas-phase basicity^46^, and aspartic acid (D) forming salt bridges^41, 49^. Interestingly, we observed that both glutamine (Q) and glutamic acid (E) had notably higher intrinsic basicities than asparagine (N) and aspartic acid (D), respectively. Both pairs of amino acids share the same functional group, but differ only in side-chain length, suggesting that the observed effect is associated with differences in conformational entropy^53^. Additionally, we discovered that amino acids with non-aromatic hydrocarbon side chains (A, V, I, L) promote overcharging. As non-polar moieties increase a peptide’s affinity for the droplet-air interface^50^, this finding suggests that peptides positioned closer to the droplet surface may exhibit a stronger overcharging response. For undercharged peptides, our results align with the effects of Coulombic repulsion for basic amino acids at the N-terminal. We also found unexpected features that counteract undercharging, such as the presence of proline, especially when located at the N-terminal, and hydrophobic amino acids (I, L, F), specifically in the timsTOF datasets. We speculate that internal prolines may reduce undercharging by introducing a kink in the peptide chain^54^, thereby promoting charge solvation^42^ and protecting from Coulombic repulsion. Further studies would be necessary to elucidate the mechanisms underlying the aforementioned observations, and why the sequence determinants differ so much in the two regimes.

Our findings suggest that peptide CSDs—through their rich sequence determinants and their differential response to experimental conditions—contain information that may benefit shotgun proteomics, information that is mostly neglected in state-of-the-art identification pipelines^55, 56^. In addition to data-dependent acquisition (DDA)^57^, our findings may especially be useful for data-independent acquisition (DIA)^58^ and MS1-only^59^ approaches, where there are no fragmentation spectra dominated by one peptide.

Of the many machine learning techniques available to analyze large-scale datasets, deep learning has had tremendous progress in recent years, as demonstrated by accurate predictions for retention time^29, 60–62^, collisional cross section^62, 63^, fragmentation spectra^31, 62^, and charge state distributions^29^. However, interpreting the predictions of black box neural networks still remains a challenge^28, 64^. While we have experimented with deep learning in the early stages of this study, we show here that classic machine learning approaches still have merit in providing easily communicable and readily interpretable insights. In fact, these two approaches are not mutually exclusive: the insights derived here from analyzing peptide CSDs can potentially inform model design and improve deep learning predictions.

Our study has several inherent limitations. Firstly, the extraction scheme cannot extract intensities that fall below the mass spectrometer’s intensity threshold. This may introduce biases in CSD readings for less abundant peptides and for scans located at the tails of elution profiles. We address this through our extraction scheme’s filtering steps which favor the extraction of high intensity peptides; we verified that the intensity threshold only accounts for < 2% error for most CSD readings (data not shown). Secondly, the distribution of peptide sequences in our dataset is affected by the proteome, the experimental setup, and the extraction scheme’s filtering steps. To address this, we used LC-MS/MS runs across different experimental parameters, proteases, cell lines, and instruments. This issue can be further addressed through incorporating synthetic peptides. Thirdly, our analysis on calculated intrinsic basicity does not capture pairwise, nor higher-order, interactions between amino acids. Although we explored including these higher-order terms, we had difficulty interpreting the results due to overparameterization. A deeper mechanistic understanding may provide opportunities for better feature engineering that can overcome these challenges. Fourthly, CSDs are influenced by all factors that occur in and downstream of ESI, including ionization efficiency, transport efficiency (from ESI to MS detection)^50^, and possible interactions within the drift tube^65^. These factors can limit the interpretation of our analysis. We partly addressed this by using runs from both Orbitrap and timsTOF instruments. Lastly, our analysis can only establish correlations, not causal relationships. Further experiments and molecular dynamics simulations are necessary to fully establish the mechanisms underlying the insights we have identified.

In conclusion, our high-throughput study of peptide CSDs has yielded informative results for the field of ESI and LC-MS/MS-based proteomics. The insights we have generated for peptide ionization contribute to our understanding of ESI mechanisms, and offer opportunities for improved peptide identification.

## Data availability

Raw files and MaxQuant analyses for our LC-MS/MS runs are available at the MassIVE data repository with ID MSV000091473. The CSD dataset generated and analyzed in this study is available at figshare^66^.

## Code availability

Extraction scheme and figure generating code are available on GitHub (https://github.com/regev-lab/extract-csd).

## Acknowledgements

We are grateful to Lars Konermann, Brian Chait, David Fenyő, and Nikolai Slavov for their insightful comments. We thank Andrew Lemoff and the UTSW Proteomics Core for their assistance with technical queries. We also thank Susan Liao for feedback on the manuscript. We acknowledge funding from Simons Investigator Award (O.R., A.M.X.); NSF MCB-2226731 (O.R.); NSF (Award 2224211) (M.J.); NIH (R35GM128802, R01AG071869 and R01HG012216) (M.J.); Columbia startup funding (M.J.); and NSF-GRFP (Award DGE2036197) (L.C.T.).

## Author contributions

A.M.X., M.J., and O.R. conceived and designed the study. L.C.T. and M.J. performed the proteomics experiments. A.M.X. processed the data and performed data analysis. A.M.X., O.R., with contributions from M.J., interpreted the results and wrote the manuscript. A.M.X. created the visualizations. O.R. supervised the project.

## Methods

### Sample preparation & mass spectrometry

To include LC-MS/MS runs with different experimental parameters, we ran our own experiments varying ESI voltage, gradient length, and sample flow rate.

HeLa cells were grown to 90% confluency and washed twice with PBS before direct lysis on the plate. Total proteins were extracted using a urea lysis buffer (8 M Urea, 75 mM NaCl, 50 mM Tris/HCl pH 8.0, 1 mM EDTA). The protein concentration was determined by Pierce BCA assay. 20 *µ*g of total protein was processed further. Disulfide bonds were reduced with 5 mM dithiothreitol and cysteines were subsequently alkylated with 10 mM iodoacetamide. Samples were diluted 1:4 with 50 mM Tris/HCl (pH 8.0) and sequencing grade modified trypsin (Promega) was added in an enzyme-to-substrate ratio of 1:50. After 16 h of digestion, samples were acidified with 1% formic acid (final concentration). Tryptic peptides were desalted on C18 StageTips according to Rappsilber et al.^67^ and evaporated to dryness in a vacuum concentrator. Desalted peptides were reconstituted in Buffer A (3% acetonitrile, 0.2% formic acid).

For mass spectrometer runs that were run in April 2021, 2 *µ*g of peptides were analyzed on a Thermo Scientific Orbitrap Q Exactive HF mass spectrometer coupled via a 25 cm long, 1.6 *µ*m particle size Aurora C18 column (IonOpticks) to an Acuity M Class UPLC system (Waters). For the long gradient, peptides were separated at a flow rate of 400 nL/min with a linear gradient spanning 2 min from 5% to 8% solvent B (100% acetonitrile, 0.1% formic acid), followed by a 87 min linear gradient from 8% to 22% solvent B, a 20 min linear gradient from 22% to 30% solvent B, a 14 min linear gradient from 30% to 60% solvent B, and a 1 min linear gradient from 60% to 90% solvent B. Each sample was run for 160 min, including sample loading and column equilibration times. For the short gradient, peptides were separated at a flow rate of 400 nL/min in linear steps from 2% to 8% solvent B over 1 min, from 8% to 30% solvent B over 33 min, from from 30% to 60% solvent B over 5 min, and from 60% to 90% solvent B over 5 min. Each sample was run for 90 min, including sample loading and column equilibration times.

For mass spectrometer runs that were run in August 2021, 2 *µ*g of peptides were analyzed on a Thermo Scientific Orbitrap Q Exactive HF mass spectrometer coupled via a 15 cm long, 3 *µ*m particle size EASY-Spray C18 column (ThermoFisher Scientific) to an Acuity M Class UPLC system (Waters). Peptides were separated at varying flow rates ranging from 200 nL/min to 800 nL/min in 200 nL/min increments. The peptides were separated in linear steps from 5% to 8% solvent B over 4 min, from 8% to 14% solvent B over 45 min, from 14% to 22% solvent B over 45 min, from 22% to 30% solvent B over 20 min, from 30% to 60% solvent B over 9 min, and from 60% to 90% solvent B over 1 min. Each sample was run for 190 min, including sample loading and column equilibration times.

Data was acquired in data dependent mode using the Xcalibur 4.1 software. MS1 spectra were measured with a resolution of 120,000, an AGC target of 3e6 and a mass range from 300 to 1800 *m*/*z*. Up to 12 MS2 spectra per duty cycle were triggered at a resolution of 15,000, an AGC target of 1e5, an isolation window of 1.6 *m*/*z* and a normalized collision energy of 28.

### Gathering published LC-MS/MS raw files

To supplement our LC-MS/MS runs, we gathered published raw files from two sources: Confetti^30^ and Meier et al.^31^. These raw files are located at the ProteomeXchange Consortium via PRIDE partner repository^68^ with data identifier PXD000900 (Confetti) and PXD019086 (Meier et al.).

### Preprocessing steps for LC-MS/MS raw files

To identify peptides that were present in the LC-MS/MS runs, the raw LC-MS/MS files were analyzed with the MaxQuant software^55^. MaxQuant was run with the following parameters: maximum of two missed cleavages, methionine oxidation and N-terminal acetylation as variable modifications, cysteine carbamidomethylation (ours, Meier et al.) or N-ethylmaleimidation (Confetti) as a fixed modification, minimum peptide length of 7 amino acids, maximum peptide mass of 4600 Da, and a false discovery rate of 1%. MaxQuant was run with version 1.6.10.43 (ours, Confetti) and version 1.6.5.0 (Meier et al., original analysis retained).

Additional preprocessing steps were performed to access MS1 profile peaks from the raw LC-MS/MS files. The Thermo .raw files from Orbitrap runs were converted to .mzML format using MSConvert from ProteoWizard version 3.0^69^, with the vendor peak picking setting enabled to obtain centroided MS1 peaks. The MS1 spectra of Bruker .d folders from timsTOF runs were accessed using alphatims v. 1.0.0^70^. The MS1 profile peaks were centroided using our own procedure (Supplementary Text 2). These centroided MS1 peaks were fed into our downstream CSD extraction scheme.

### CSD extraction scheme

Per-scan CSD readings were extracted from MS1 scans for MS2-identified peptides using the following scheme. For each peptide and each charge state from 1^+^ to 5^+^, relevant MS1 scans were searched for peaks that matched the theoretical isotope envelope and passed stringent filtering requirements. From these peaks, the charge states’ intensities were estimated, and then normalized to obtain the peptide’s CSD reading for that scan. Finally, these per-scan CSDs were combined into one CSD estimate per peptide by performing an intensity-weighted average across scans. The full details of the extraction scheme are provided in Supplementary Text 2.

One of the main design goals was to achieve high quality CSD readings. As such, we applied stringent filtering and only retained CSD readings which contained confident intensity estimates for every charge state. To determine whether a peptide’s charge state was present in an MS1 spectrum, we verified that (i) the theoretical isotope peaks had low *m*/*z* offset from the observed isotope peaks, (ii) the shape of the theoretical isotope distribution matched the observed isotope distribution (cosine similarity >0.98), and (iii) there was an absence of peaks that might belong to the isotope distributions of other peptides (chimeric peaks). On the other hand, a charge state was denoted absent and thereby assigned an extracted intensity of 0 if no peaks were in a sufficiently large *m*/*z* vicinity. Through tuning the thresholds used for filtering, the extraction scheme favors the extraction of highly confident peptide CSD readings.

We analyzed a total of 326 raw LC-MS/MS files which resulted in CSD readings extracted for a total of 261,667 unique peptides (Table 1). The resulting CSD dataset has a similar distribution of peptides to those typically identified using MS/MS, with masses ranging from 700 Da–4600 Da, and varying numbers of basic sites (Fig. 2).

### Measuring error between CSD readings

Error between CSD readings of the same peptide was measured through total variation. The total variation between two distributions (*p*_1_, . . . , *p_n_*) and (*q*_1_, . . . , *q_n_*) is given by the sum of absolute differences between probabilities divided by two:

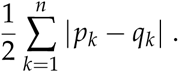

Total variation ranges from 0 to 1, signifying equivalent or disjoint distributions, respectively.

### Correcting experimental batch effects

To better assess CSD errors across pairs of runs, we applied a one-parameter batch correction (Supplementary Figs. 1,2). Peptide CSDs were transformed according to the following scheme. For each charge state *k*^+^, we scaled its probability *p_k_*_+_ by a factor of *β^k^*, for some global batch parameter *β* > 0. The scaled probabilities were then renormalized to sum to one, forming the transformed CSD. In other words, our batch correction maps peptide CSDs of the form (*p*_1+_ , . . . , *p*_5+_) to

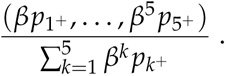

This transformation can be interpreted as additive shifts in the energy scale (i.e., log probability scale), where the magnitude of the shift scales linearly with the charge state.

Errors across pairs of runs were measured before and after applying a batch correction (Supplementary Fig. 1). The batch correction parameter was chosen to minimize the total error, calculated as the sum of the total variation in CSD readings across all peptides common to both runs. We found this batch correction to be reasonably effective given that it only depends on one parameter.

### Calculating and analyzing intrinsic basicity scores

In the analysis of sequence effects on charging, we calculated the intrinsic basicities of amino acid features for each of four charging regions (Fig. 3). Each charging region considered a subset of peptides (those with a given number of basic sites and with no variable modifications) and charging across consecutive charge state probabilities *p_k_*_+_ and *p*_(*k*+1)+_ for some *k* from 2 to 4.

The calculated intrinsic basicities were given by the coefficients of a logistic regression. The input variables were chosen as follows: 20 numerical variables for each of the 20 amino acid counts, and 20 binary variables to denote the identity of the N-terminal amino acid. The target variable for the regression was chosen as *p*_(*k*+1)+_ /(*p_k_*_+_ + *p*_(*k*+1)+_ ). The logistic regression was performed using the pyGAM library^71^, with binary cross entropy as the loss function and mild L2 regularization (*λ* = 0.01).

To compare across region-run pairs (Fig. 3c), we calculated the Pearson correlation coefficient of the calculated intrinsic basicities for the 17 non-basic amino acids. Only our LC-MS/MS runs with sufficiently many data points in all four charging regions were considered; that is, each charging region needed to contain >100 peptides that exhibited nonzero probabilities on both charge states in question. The resulting LC-MS/MS runs, which were used in Fig. 3c, are the 12 runs from the April 2021 experiment.

To calculate mass-adjusted intrinsic basicities for each region-run pair (Fig. 3d), we subtracted from the overall intrinsic basicities the portion that could be explained by mass. Specifically, the mass-adjusted intrinsic basicities were derived as the residuals of a linear regression between the calculated intrinsic basicities and the masses of the 17 non-basic amino acids plus an intercept term. The linear regression used a Huber loss^72^ with parameter *δ* = 0.01. We selected the Huber loss, which is equal to the L1 loss for large values (>*δ*) and the L2 loss for small values (<*δ*), to ensure that amino acids with intrinsic basicities significantly different from the mass trend were not overly penalized, and to guarantee uniqueness of the coefficients.

## Extended Data Figures

**Supplementary Figure 1.**
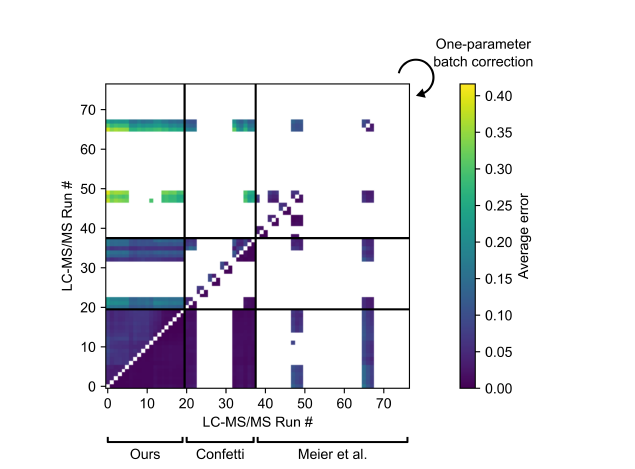
Error across runs before and after batch correction. Color-coded matrix of average CSD error across pairs of runs (with *≥* 200 common peptides) measured before (upper left) and after (lower right) applying a one-parameter batch correction (see methods). Error is measured as average total variation (see methods) between CSD readings of common peptides. Meier et al.’s runs were aggregated across fractionations (average taken for duplicate readings).

**Supplementary Figure 2.**
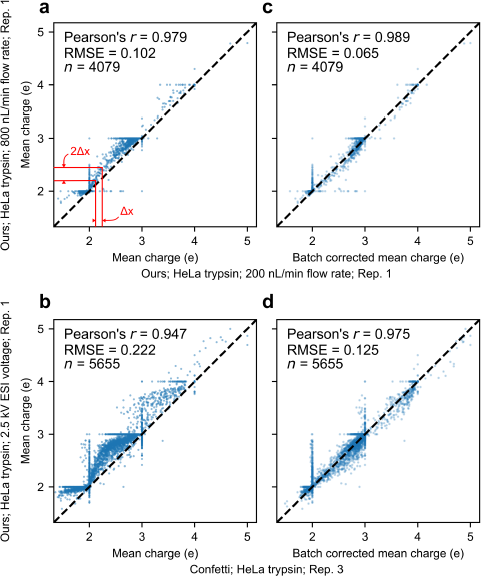
Examples of before and after batch correction. Comparison of mean charge readings of common peptides between (a,c) our low and high flow rate runs, and (b,d) a Confetti run and our high voltage run. Mean charge is shown before (a,b) and after (c,d) applying a one-parameter batch correction (see methods). Red lines and text in (a) illustrate the effect of experimental variation on mean charge differences.

**Supplementary Figure 3.**
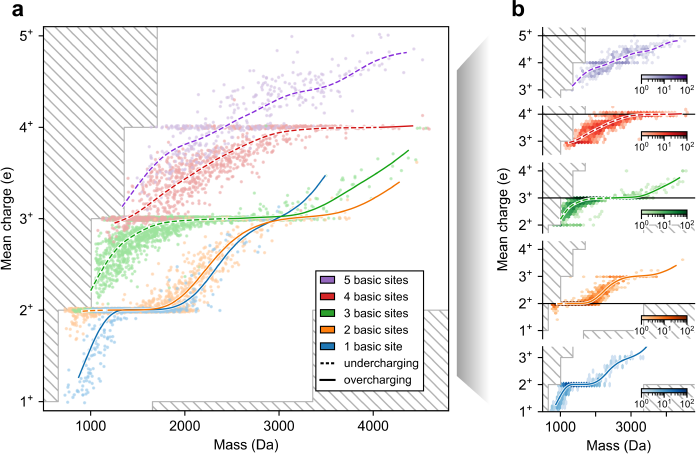
Visualizing CSD dataset versus mass for a Confetti run. Plots of mean charge versus mass for peptides from a representative run (Confetti; HeLa GluC) as in Fig. 2.

**Supplementary Figure 4.**
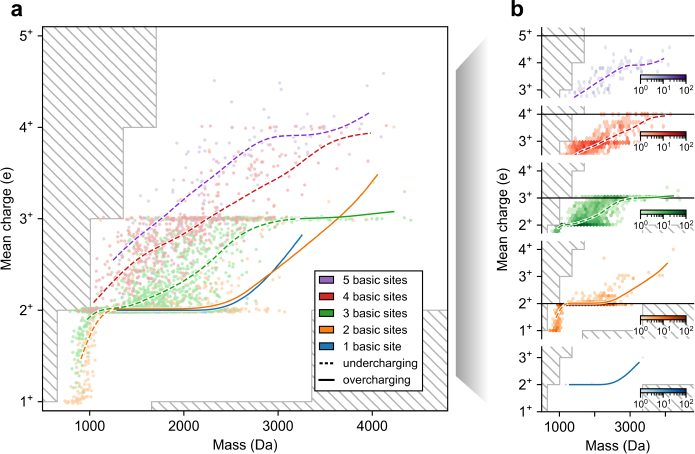
Visualizing CSD dataset versus mass for a run from Meier et al. Plots of mean charge versus mass for peptides from a representative run (Meier et al.; HeLa trypsin) as in Fig. 2

**Supplementary Figure 5.**
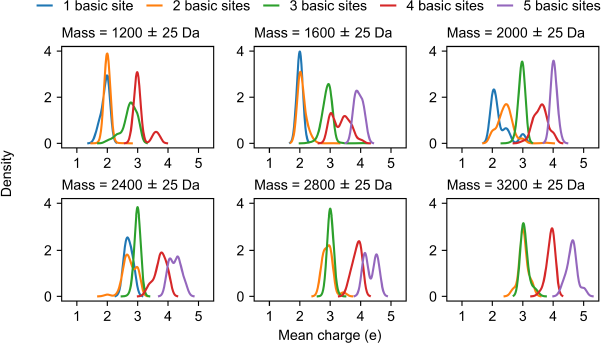
Mean charge for different basic site counts are generally disjoint. Distributions of peptide mean charge, separated by number of basic sites, shown along different slices of mass. Kernel density estimation is performed with a gaussian kernel (bandwidth = 0.1). Data shown is from our HeLa trypsin run with 2.5 kV ESI voltage, 160 min gradient length, 400 nL/min flow rate.

**Supplementary Figure 6.**
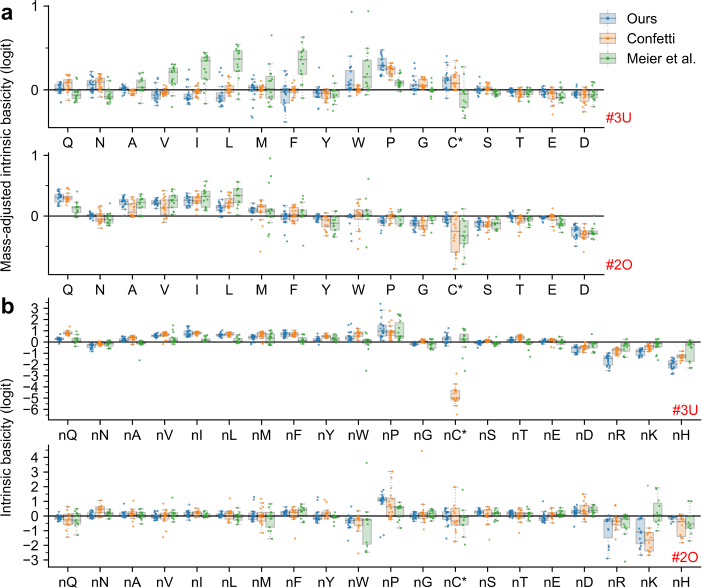
Extended versions of. Figure 3d,e. Box plot (with outliers) of (a) estimated mass-adjusted intrinsic basicities (given by residual intrinsic basicities after subtracting trend in mass, see methods) for amino acids and (a) intrinsic basicities for identity of N-terminal across runs, separated by data source and charging region. C* denotes carbamidomethyl-cysteine in our and Meier et al.’s runs, and N-ethylmaleimide modified cysteine in Confetti’s runs. Box plot elements: centerline, median; boxplot limits, upper and lower quartiles; whiskers, 1.5x interquartile range; points, all with jitters.

**Supplementary Figure 7.**
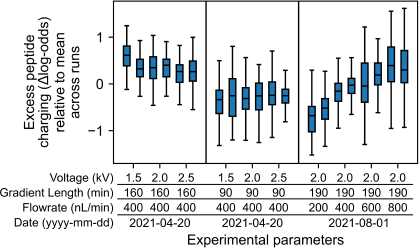
Relative charging across runs. Box plot showing distribution of excess charging across runs for 124 peptides common to all our runs. Excess peptide charging is the difference between a peptide’s mean log odds across its elution to the mean of that value across all runs. Log odds is defined as the log of the ratio of consecutive charge states. Box plot elements: centerline, median; boxplot limits, upper and lower quartiles; whiskers, 1.5x interquartile range; points, none (outliers not shown).

**Supplementary Figure 8:**
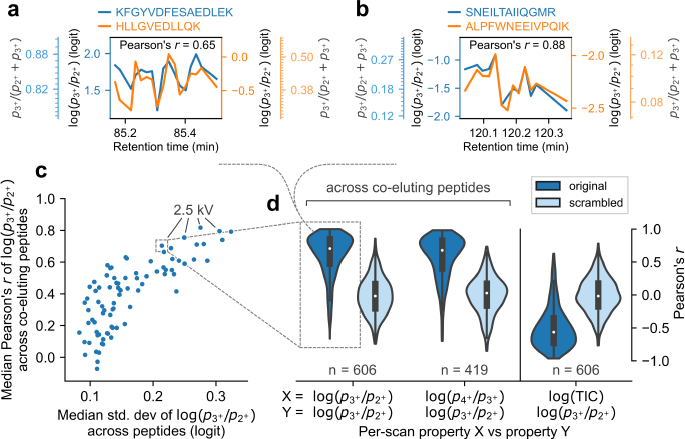
Per-scan CSD fluctuations and effect of experimental parameters on charging. (a,b) Fluctuations in CSD readings of two representative pairs of co-eluting peptides across their common scans. (c) Scatterplot, across runs, of median correlation between CSD fluctuations of co-eluting peptides (*>*10 common scans) versus median standard deviation in CSD fluctuation. Runs were only included if they contained *>*15 co-eluting peptides. Our runs with 2.5 kV ESI voltage are labeled. CSD fluctuations in (a–c) are measured as the log odds between charge state 3^+^ and 2^+^. (d) Violin plot showing correlations between log odds of co-eluting peptide CSDs (left and middle columns), and between log odds of peptide CSDs and total ion current (right column) for our HeLa trypsin run (2.5 kV ESI voltage, 160 min gradient length, 400 nL/min flow rate). Correlations taken after scrambling CSD readings across co-elution are shown as controls. Violin plot elements identical to previously defined box plot elements.

**Supplementary Figure 9.**
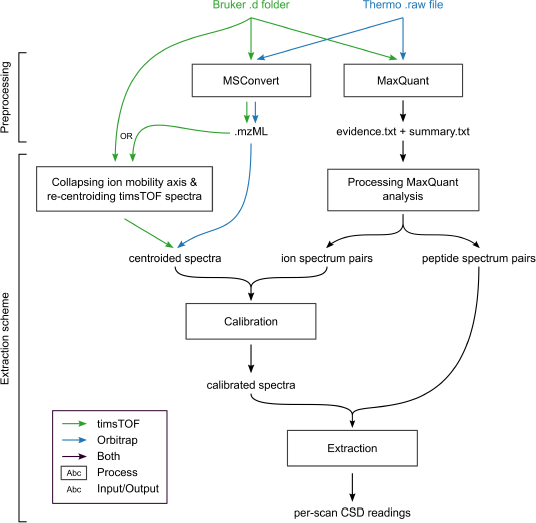
Infographic for the extraction scheme. Flowchart outlining steps taken to process raw LC-MS/MS files to obtain per-scan CSD readings.

## Supplementary Text

### Supplementary Text 1: CSD variations across scans offer opportunities for identification

In the main text, we assigned a single CSD reading to each peptide, which was obtained through averaging per-scan CSD readings across the peptide’s elution. In this section, we report our findings on how CSD readings for the same peptide differ across scans.

We observed that CSDs exhibited mild scan-to-scan fluctuations, ranging from *∼*1–5% depending on the LC-MS/MS run. To verify whether these CSD fluctuations were due to noise or due to scan-to-scan differences in overall charging, we compared the fluctuations of co-eluting peptides (Supplementary Fig. 8a,b). In most runs, we found that co-eluting peptides exhibited high correlation in their scan-to-scan charging, the only exceptions being runs that showed little to no fluctuations (Supplementary Fig. 8c,d). Moreover, a representative high-correlation run (one of our 2.5 kV runs) demonstrates that strong correlation is present not only on average but also for nearly all pairs of co-eluting peptides (Supplementary Fig. 8d, left). These correlations were also consistently observed regardless of the choice of charge states: comparing charging across 2^+^ and 3^+^ from one peptide with charging across 3^+^ and 4^+^ from another co-eluting peptide showed equally strong correlations (Supplementary Fig. 8d, center). In summary, these findings indicate that CSD fluctuations are largely attributable to variations in the overall degree of charging across scans.

Next, we sought to identify per-scan features that correlated with CSD fluctuations and can provide insights into their underlying cause. Among the features available from the LC-MS/MS files, we identified negative correlation (median Pearson’s *r* across peptides = *−*0.57) between per-scan charging and total ion current (TIC), defined as the total intensity of all ion peaks in the scan (Supplementary Fig. 8d, right). Since ion current is proportional to charge times molar abundance, the observed negative correlation between charging and TIC further indicates a negative correlation between charging and total molar abundance of analytes entering the mass analyzer. One explanation for this observation is the phenomenon of “charge competition”: prior work has demonstrated that higher charge availability per analyte (for example from decreasing analyte concentration) resulted in higher charging^35, 73^. As such, these results suggest that fluctuations in the total abundance of analytes (against a fixed background of charge availability) cause changes in scan-to-scan peptide charging due to fluctuating levels of “charge competition”. Overall, these finding indicate that correlated CSD fluctuations arise from some underlying, time-varying experimental factors.

These findings have potential applications to mass spectrometry identification. Specifically, peptides that have similar CSDs in one scan may exhibit more disparate CSDs in another scan due to differences in charging conditions, similar to what was observed for run-to-run CSD variations (see main text). For example, consider the two co-eluting peptides SNEILTAIIQGMR and ALPFWNEEIVPQIK during the 13 scans shown in Supplementary Fig. 8b. During the low charging scans, the absolute differences in CSD is *∼* 7%, whereas during the high charging scans that differences is *∼* 14%. Namely, this proof-of-concept example showcases that scan-to-scan charging variations may amplify differences in CSDs, allowing them to be more easily distinguished. Overall, these findings highlight potential avenues for enhancing peptide identification through leveraging scan-to-scan CSD fluctuations.

### Supplementary Text 2: Details of CSD extraction scheme

Here, we provide details about the CSD extraction scheme (see infographic, Supplementary Fig. 9). In this section, the terms “scans” and “spectra” both refer to MS1 mass spectra (at a specific retention time). The term “ion” refers to a peptide combined with a charge state. Adjustable parameters of the extraction scheme are denoted in all caps (default values are provided in “Default parameters” below).

### Collapsing ion mobility axis & re-centroiding timsTOF spectra

*Input:* Bruker .d folder or .mzML file

*Output:* centroided spectra (without ion mobility axis)

Here, we describe the steps taken to collapse the ion mobility axis, and re-centroid the resulting peaks for timsTOF MS1 spectra. First, we collapsed the ion mobility axis, retaining the *m*/*z*, and intensity axes. Second, we filtered all peaks with intensity lower than CENTROID-ING MIN PROFILE PEAK INTENSITY. Third, we identified peaks that were adjacent in *m*/*z* (where two peaks are considered adjacent if they are located consecutively based on the uniform discretization in the “time of flight” scale, that is in the (*m*/*z*)^2^ scale). Third, we grouped peaks based on adjacency (taking the maximum range of peaks that were adjacent to one another). Fourth, for each group of peaks, we assigned one centroided peak, with intensity equal to the sum of intensities and *m*/*z* equal to the intensity-weighted average of *m*/*z*’s.

### Processing MaxQuant analysis

*Input:* MaxQuant analysis (evidence.txt, summary.txt)

*Output:* list of ion spectrum pairs, list of peptide spectrum pairs

Here, we describe the steps taken to establish ground truth ions and peptides, and the spectra they are located in, which will be used in the later calibration and extraction stages.

For each identified ion from the MaxQuant analysis, we extracted the start and finish elution times based on those stated in the evidence.txt file. Since each ion may appear multiple times in the evidence.txt file, the start (finish) time was defined as the minimum (maximum) of all occurrences of that ion.

For each identified ion, we paired the ion with all spectra that had a retention time located between the ion’s start and finish elution time. For the purposes of quality control, an ion was removed if the start and finish time differed by more than CALIBRATION MAX RETENTION LENGTH.

For each identified peptide, we paired the peptide with all spectra that were previously paired to one of the peptide’s ions. In other words, the peptide spectrum pairs are a union of its ion spectrum pairs.

### Calibration

*Input:* centroided spectra

*Output:* calibrated spectra

To improve the subsequent extraction stage, we calibrated the *m*/*z* axis of the MS1 spectra using the following steps. First, for each ion spectrum pair (see “Processing MaxQuant analysis” above), we computed its *m*/*z* offset, given as the difference between the ion’s theoretical monoisotopic *m*/*z* and its nearest peak in the spectrum. Then, for each spectrum, we calculated a robust average (see below) of all the *m*/*z* offsets of paired ions. Lastly, we shifted the *m*/*z* of all peaks in the spectrum by that average.

To compute the robust average of *m*/*z* offsets, we first computed a rough estimate for the FWHM (full width at half maximum), calculated as 2 median(*|m*/*z* offsets *−* median(*m*/*z* offsets)*|*). Then, we removed all *m*/*z* offsets with magnitude greater than 3 FWHM. Lastly, we took the average of the remaining *m*/*z* offsets.

To avoid problematic scans, spectra with an insufficient number of ions were removed from downstream steps, where the minimum number of required ions is determined by CALIBRATION MINIMUM REQUIRED PEPTIDE COUNT.

### Extraction

*Input:* calibrated spectra

*Output:* per-scan CSD readings

Here, we describe the steps taken to extract per-scan CSD readings for each peptide spectrum pair (see “Processing MaxQuant analysis” above). As an overview, peptide CSDs were calculated through normalizing the estimated intensity readings of each charge state. To ensure high-quality extractions, charge state intensity readings were labeled as “confidently present”, “confidently absent”, or “ambiguous”, and CSDs were filtered if they contained any “ambiguous” intensity readings. Below, we describe the specific details of the extraction scheme and the labeling criteria used.

For each peptide spectrum pair, we first computed properties regarding the theoretical and observed isotope distributions. For each charge state from 1^+^ to 5^+^ (or EXTRACTION MAX CHARGE), we computed the first three peaks of the theoretical isotope distribution; we refer these three peaks as #0 (monoisotopic), #1, and #2 peaks, respectively. We also extracted the isotope distribution observed in the spectrum, defined as the collection of the nearest peaks (in *m*/*z*) to the theoretical #0, #1, and #2 peaks. From these, we calculated the following properties: the *m*/*z* offsets between the theoretical and observed isotope peaks, the cosine similarity between the isotope distributions, and dot product between the isotope distributions (which serves as the charge state’s estimated intensity reading). Moreover, we checked for the presence of extraneous peaks that may suggest that the observed isotope distribution overlaps with other peptide spectra. Namely, we extracted the nearest peak to the theoretical #-1 peak (a peak located one neutron below the monoisotopic peak). We also extracted nearest peaks to the theoretical #-1/2, #1/2, and #3/2 peaks (which are located at the midpoints of the #-1, #0, #1, and #2 peaks). Moreover, these extraneous peaks were denoted as non-negligible if their intensities were high (defined as greater than one half of the intensities of adjacent #0, #1, #2 peaks). Lastly, these properties were used to label the given charge state based on the criteria below.

Charge states were labeled as “confidently present” if all three conditions held:

- absolute *m*/*z* offset for #0, #1, and #2 peaks < EXTRACTION MAXIMUM MZ OFFSET FOR MATCH
- absolute *m*/*z* offsets for #-1, #-1/2, #1/2, and #3/2 peaks with non-negligible intensities > EXTRACTION MAXIMUM MZ OFFSET FOR EXTRANEOUS PEAKS
- cosine similarity score > EXTRACTION MINIMUM SIMILARITY SCORE FOR MATCH

and consequently assigned an intensity reading derived from the spectrum (see above). Charge states were labeled as “confidently absent” if the following condition held:

- absolute *m*/*z* offset for #0 and #1 > EXTRACTION MINIMUM MZ OFFSET FOR NO MATCH

and consequently assigned an intensity reading of zero. Charge states were labeled as “ambiguous” otherwise.

If none of the charge states were labeled as “ambiguous”, then the peptide was assigned a CSD reading for the scan, given by the normalized intensity readings. For the results in the main text, each peptide was assigned a single CSD reading through an intensity-weighted average of CSDs across scans.

### Default parameters

The default values for the parameters of the extraction scheme are:

- CENTROIDING MIN PROFILE PEAK INTENSITY = 100
- CENTROIDING MIN CENTROIDED PEAK INTENSITY = 500
- CALIBRATION MAX ABS MZ OFFSET = 0.05
- CALIBRATION MAX RETENTION LENGTH = 5.0
- CALIBRATION MINIMUM REQUIRED PEPTIDE COUNT = 5
- EXTRACTION MAX CHARGE = 5
- EXTRACTION MINIMUM SIMILARITY SCORE FOR MATCH = 0.98
- EXTRACTION MAXIMUM MZ OFFSET FOR MATCH = 1×FWHM
- EXTRACTION MAXIMUM MZ OFFSET FOR EXTRANEOUS PEAKS = 2.5×FWHM
- EXTRACTION MINIMUM MZ OFFSET FOR NO MATCH = 5×FWHM

where FWHM is determined from the absolute *m*/*z* offsets between the theoretical and observed monoisotopic peaks in the “Extraction” stage (e.g., 0.0018 Da for our HeLa trypsin run with 2.5 kV ESI voltage, 160 min gradient length, 400 nL/min flow rate).

